# Investigating CRISPR spacer targets and their impact on genomic diversification of *Streptococcus mutans*

**DOI:** 10.1101/2022.04.14.488371

**Authors:** Alejandro R. Walker, Robert C. Shields

## Abstract

CRISPR-Cas is a bacterial immune system that restricts the acquisition of mobile DNA elements. These systems provide immunity against foreign DNA by encoding CRISPR spacers that help target DNA if it re-enters the cell. In this way, CRISPR spacers are a type of molecular tape recorder of foreign DNA encountered by the host microorganism. Here, we extracted ∼8,000 CRISPR spacers from a collection of over three hundred *Streptococcus mutans* genomes. Phage DNA is a major target of *S. mutans* spacers. Strains have also generated immunity against mobile DNA elements such as plasmids and integrative and conjugative elements. There may also be considerable immunity generated against bacterial DNA, although the relative contribution of self-targeting versus bona fide intra- or inter-species targeting needs to be investigated further. While there was clear evidence that these systems have acquired immunity against foreign DNA, there appeared to be minimal impact on horizontal gene transfer (HGT) constraints on a species-level. There was little or no impact on genome size, GC content and ‘openness’ of the pangenome when comparing between *S. mutans* strains with low or high CRISPR spacer loads. In summary, while there is evidence of CRISPR spacer acquisition against self and foreign DNA, CRISPR-Cas does not act as a barrier on the expansion of the *S. mutans* accessory genome.

**Impact Statement:** CRISPR-Cas is a widespread bacterial immune system that has been repurposed as a molecular biology tool. This study investigates the role of these systems in the biology and evolution of the dental caries pathogen *Streptococcus mutans*. CRISPR spacers, that encode immunity against foreign DNA, were extracted from over three hundred *S. mutans* isolates. Sequence analysis showed that the CRISPR spacers match against phage, mobile element, and bacterial DNA. This shows that *S. mutans* is actively acquiring immunity against horizontally acquired DNA. However, additional analysis revealed little to no impact of CRISPR-Cas systems on diversification of the *S. mutans* genome. This suggests that while these systems are actively acquiring CRISPR spacers to defend against foreign DNA, the overall impact on the *S. mutans* genome might be small.

**Data Summary:** Supporting data provided on the Github platform: https://github.com/theshieldslab/Streptococcus-mutans-CRISPR-Spacers-Analysis

The authors confirm all supporting data, code and protocols have been provided within the article or through supplementary data files.

## Introduction

*Streptococcus mutans* is a Gram positive bacteria that inhabits the oral cavity and has a strong association with causing dental caries (tooth decay) [1]. It has a genome size of approximately 2 Mbp and an “open” pangenome, meaning that as more *S. mutans* genomes are sequenced new genes previously not associated with *S. mutans* are discovered [2, 3]. These observations suggest that the organism is adaptable and able to acquire new traits over time. More broadly, it has been estimated that as a group 60%+ of the pangenome of streptococci is acquired via horizontal gene transfer (HGT) [4]. The ability of *S. mutans* to horizontally acquire DNA is likely greatly facilitated by competence development, a trait that is conserved (*comRS, comX* and late-competence genes) across the species [5]. There are other important contributors to the genetic variability observed in *S. mutans* including residing in the oral microbiome which is likely a hotspot for HGT and significant pressure to adapt to new eco-niches. Regarding the last point, *S. mutans* strains have acquired traits, such as collagen-binding proteins, that have expanded its niche into cardiovascular diseases [6].

In a recent publication we detected the presence of at least one CRISPR (clustered regularly interspaced short palindromic repeats)-Cas(CRISPR-associated protein) system in 95% of *S. mutans* strains from a collection of 477 genomes [7]. CRISPR-Cas systems function as an adaptive immune system, incorporating DNA fragments from invading DNA (CRISPR spacers) that in concert with Cas endonucleases prevent future ‘re-infection’ of the invading DNA. CRISPR-Cas systems are organized into types and subtypes depending on the function of the Cas nuclease and the organization of the CRISPR-Cas loci. Type I (C and E subtypes) and Type II (A subtype) are most commonly identified in *S. mutans* and these systems recognize and cleave DNA. There are rare instances where Type III-A systems, that have specificity towards cleaving RNA [8], are found in *S. mutans*. A substantial number of *S. mutans* strains encode both a Type I-C and a Type II-A CRISPR-Cas system, perhaps to help with viral escape from either system. The model *S. mutans* strain, UA159, carries Type II-A and Type I-C CRISPR-Cas loci. The Type II-A system is functional, is capable of interfering with plasmid transformation and has been repurposed for CRISPR interference and genome editing [7, 9, 10]. It is likely that the Type I-C system is not functional because of a truncated Cas1 protein [10, 11]. Interference of both plasmid and phage DNA has also been shown in *S. mutans* P42S, with the Cas9 protein recognizing a different protospacer adjacent motif (PAM) domain compared with the UA159 Cas9 nuclease [12]. Some progress has been made exploring the functionality of these systems but there has been limited analysis of CRISPR spacers acquired by *S. mutans*. The most comprehensive study of *S. mutans* CRISPR spacers was conducted in 2009 from 29 individual strains [11]. With our large collection of *S. mutans* genomes, and changes in CRISPR-Cas analysis over the last decade, we believe an updated analysis of CRISPR spacers is warranted.

The observation that *S. mutans* has a large pan-genome is in conflict with the high frequency of CRISPR-Cas systems across the species. These systems have primarily been considered as a phage defense mechanism. However, there is experimental evidence that CRISPR-Cas systems can limit other types of HGT (*e*.*g*. plasmid acquisition) in laboratory settings including in *S. mutans* [10]. Despite evidence that CRISPR-Cas systems protect bacteria from phage and other mobile genetic element infections in simplified systems, it remains unclear how much they contribute to the ecology and evolution of bacteria living in complex natural environments. When large genome datasets are studied there has been conflicting evidence of the impact of CRISPR-Cas on HGT in bacterial populations. In *Pseudomonas aeruginosa* CRISPR-Cas systems constrain HGT [13] and there is evidence of immunity against antibiotic resistance genes in *Enterococcus faecalis* strains carrying CRISPR-Cas [14]. However, there is also evidence of no inhibition of HGT by CRISPR-Cas in other species [15]. The oral microbiome, specifically dental plaque, has a high CRISPR spacer load compared with other body sites [16], it is a dynamic environment and it is a large community of interacting microbes and phages. It therefore provides an excellent model for understanding the contribution of CRISPR-Cas in limiting HGT in complex microbial settings. Here, using *S. mutans* as a starting point, we will begin to address how impactful CRISPR-Cas is on genomic diversification in this complex ecosystem.

## Methods

### Genomic data

We have a collection of 477 *S. mutans* genomes that was curated by Vince Richards at Clemson University. Genomes are available in .gbk format on the Github platform (https://github.com/theshieldslab/Streptococcus-mutans-CRISPR-Spacers-Analysis). For this study CRISPR spacers were isolated from 335 *S. mutans* genomes. Incomplete assemblies were evident in the other *S. mutans* genomes, likely because of Illumina based whole genome sequencing and subsequent difficulty identifying repeat regions (*i*.*e*. CRISPR spacer regions).

### CRISPR spacer identification

Identification and data curation of the CRISPR systems found in our clinical isolates was conducted at the University of Florida HiPerGator cluster computer with the CRISPRCasFinder [17] version 4.2.17 pipeline. The resulting output files, containing the predicted spacers, were then processed and summarized in several files containing CRISPR system types and subtype, list of genes, repeat sequences, count of repeat sequences, and spacers sequences. The spacer sequences were then labeled with their respective isolate name, contig ID where the spacers was predicted, and an additional counter number. All CRISPR spacer sequences were then parsed into a FASTA format file, which is available on the Github platform.

### CRISPR spacer target identification

Blastn was used to identify spacer targets from phage, integrative and conjugative elements (ICEs), plasmids, and *S. mutans*. Phage DNA sequences were downloaded from NCBI (ftp://ftp.ncbi.nih.gov/refseq/release/viral/). ICE sequences were downloaded from the ICE database ICEberg 2.0 (https://bioinfo-mml.sjtu.edu.cn/ICEberg2/download.html) [18]. Plasmid sequences were downloaded from a database containing 10,892 complete and annotated plasmids [19]. In addition, known *S. mutans* plasmid sequences were compiled (available on the Github platform) [20–22]. Parameters for Blastn were set to 95% sequence coverage (qcovs) and 85% sequence identity (pident). Other parameters generated with the sequence results included the number of mismatches, the evalue and the subject strand (sstrand). All raw output data is available as .xlsx files on the Github platform.

### Analysis of genome size, GC content and the pangenome

The data mining for gene clusters was conducted in a series of steps, starting with Prokka annotations for each clinical isolate genome [23]. Since all isolates belong to the *Streptococcus* genus, annotations with Prokka were defined within Bacteria and Streptococci for better gene predictions. Multilocus sequence typing (MLST) was conducted with Roary [24], which is ideal for close non-divergent genomes. The final step was to generate the phylogenomic tree from the genome alignment file created by Roary. A maximum likelihood tree was drawn with the Phyml [25, 26] algorithm and plotted in R [27] with the GGtree [28] package. This work also included custom bash command lines in order to generate accurate measurements of GC content and genome size for all the isolates in our bank.

## Results and Discussion

### CRISPR arrays and spacers are frequently detected in *S. mutans* isolates

Using CRISPRCasFinder [17] across our genome database we identified a total of 8,078 CRISPR spacers of which 5,875 are unique sequences. Isolates with CRISPR arrays present carry an average of 23.9 spacers each, with strain smu174 carrying 129 spacers in total (on one array). By comparison, *Streptococcus thermophilus* strains carry an average of 33 spacers per cassette [29]. The majority of *S. mutans* strains carry one or two CRISPR arrays but there are rare instances of strains carrying 4+ (Fig. 1A). Figure 1B shows a frequency distribution of CRISPR spacer load across all the isolates. CRISPR spacer acquisition is a balance between obtaining immunity against harmful sequences without diluting spacers (some which may provide inadequate immunity because of mutation) across the available Cas machinery [30]. Most CRISPR spacers were either 30-nt or 32-35-nt in length (Fig. 1C), with these two groups likely arising from differences in Type I and Type II CRISPR spacer lengths. The average GC content of CRISPR spacers was 38.16%, which is above the average GC content of *S. mutans* strains (Fig. 1D). This may indicate that foreign DNA encountered by *S. mutans* is on average of a higher GC content than the average *S. mutans* genome. By way of comparison, M102AD, an *S. mutans* infecting phage has GC content of 39.6%.

**Figure 1.**
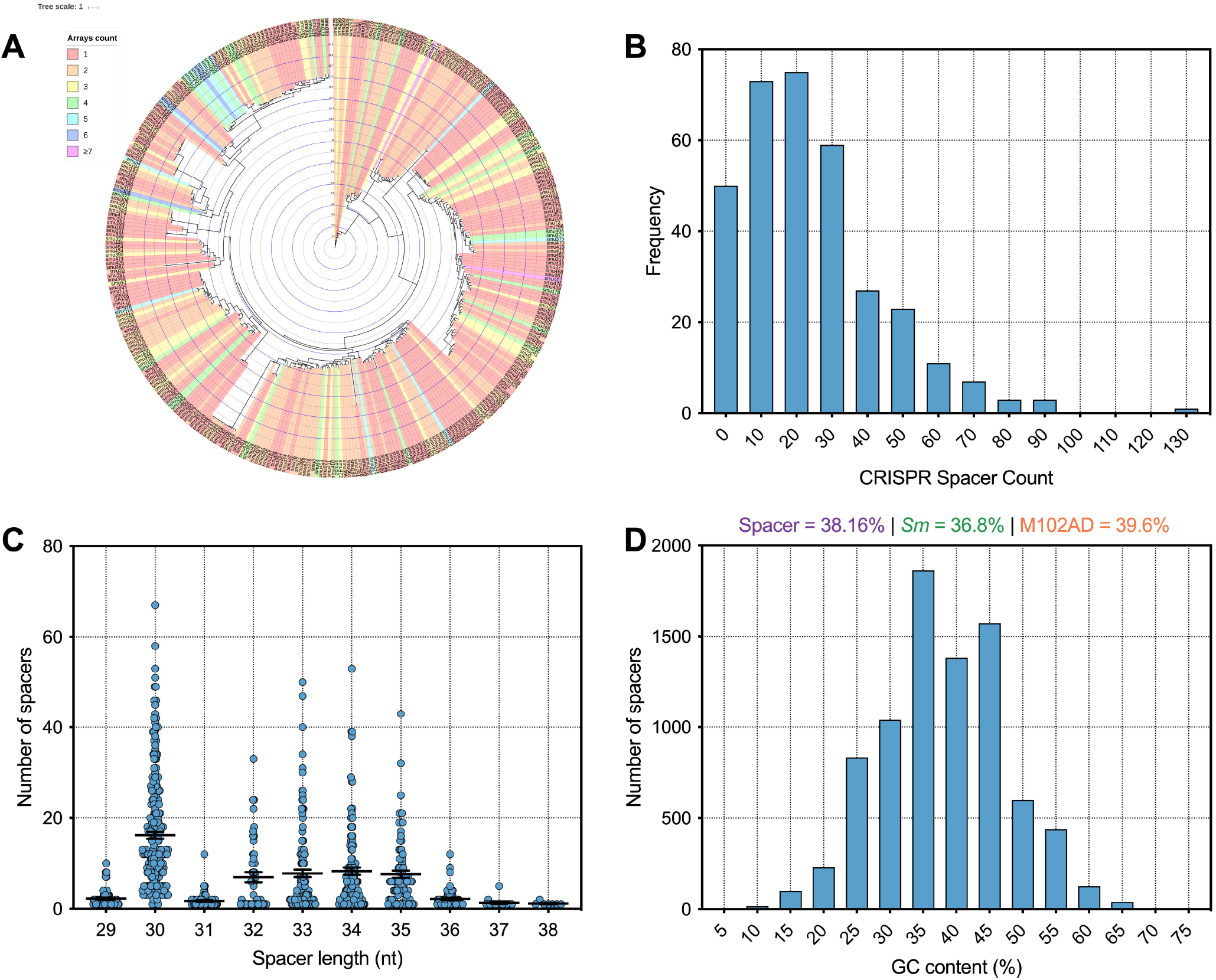
Analysis of *S. mutans* CRISPR spacers. (A) A phylogenetic tree was generated and strains were colored according to how many CRISPR spacer arrays were present on each strain genome. (B) A frequency distribution of CRISPR spacer load across all the isolates. (C) CRISPR spacers were grouped according to the length (nt) of the CRISPR spacers. (D) A frequency distribution of CRISPR spacer GC content (percentage GC) was generated.

### Targets of *S. mutans* CRISPR spacers

Having identified 8,078 spacers across our *S. mutans* genomes we next wanted to identify targets for these spacers. First, we characterized how many of these spacers target foreign DNA. This is the traditional target of CRISPR-Cas systems, and includes interference with phage, plasmid, integrative and conjugative elements (ICEs), and other bacterial DNA. In total, 1913 spacers (23.7%) mapped to these targets. Next, we will describe these targets in more detail for each sub-group of foreign DNA.

#### Phage

Adaptive immunity against bacteriophage is a notable feature of CRISPR-Cas systems across bacteria. Consistent with this concept *S. mutans* CRISPR spacers mapped to 1,137 phage sequences, of which 531 were unique matches. A total of sixteen distinct phage species are targeted by *S. mutans* CRISPR spacers (Table S1). For four of these phages *S. mutans* is the known host, whereas for the other phages the hosts are other Firmicutes bacteria; Staphylococci, Enterococci, Lactobacilli and Streptococci. For these phages, typically only one spacer aligned against a conserved target. We anticipate that rather than infecting *S. mutans*, these phage genes are present either as prophage elements or as orphan genes in *S. mutans* genomes. The four *S. mutans* infecting phage that *S. mutans* has acquired spacer immunity against are known as M102, M102AD, □APCM01 and smHBZ8 [31–34]. These are the only *S. mutans* phage that have been genome sequenced and all belong to the *Siphoviridae* family of double-stranded DNA viruses. At least two other *S. mutans* phage, e10 and f1, are known to infect *S. mutans* but these strains have not been sequenced [32]. The four sequenced *S. mutans* phage are highly related, particularly the first 20 Kbp (Figure 2). However, they each contain a variable region consisting of small open reading frames with unknown functions at the 3’ end of the genome (Figure 2). Spacers were mapped and viewed with a genome browser to determine if there is any bias in spacer acquisition to specific phage genes. Overall, it appears that immunity to the variable region is less commonly acquired, and this was most evident for M102 and M102AD.

**Figure 2.**
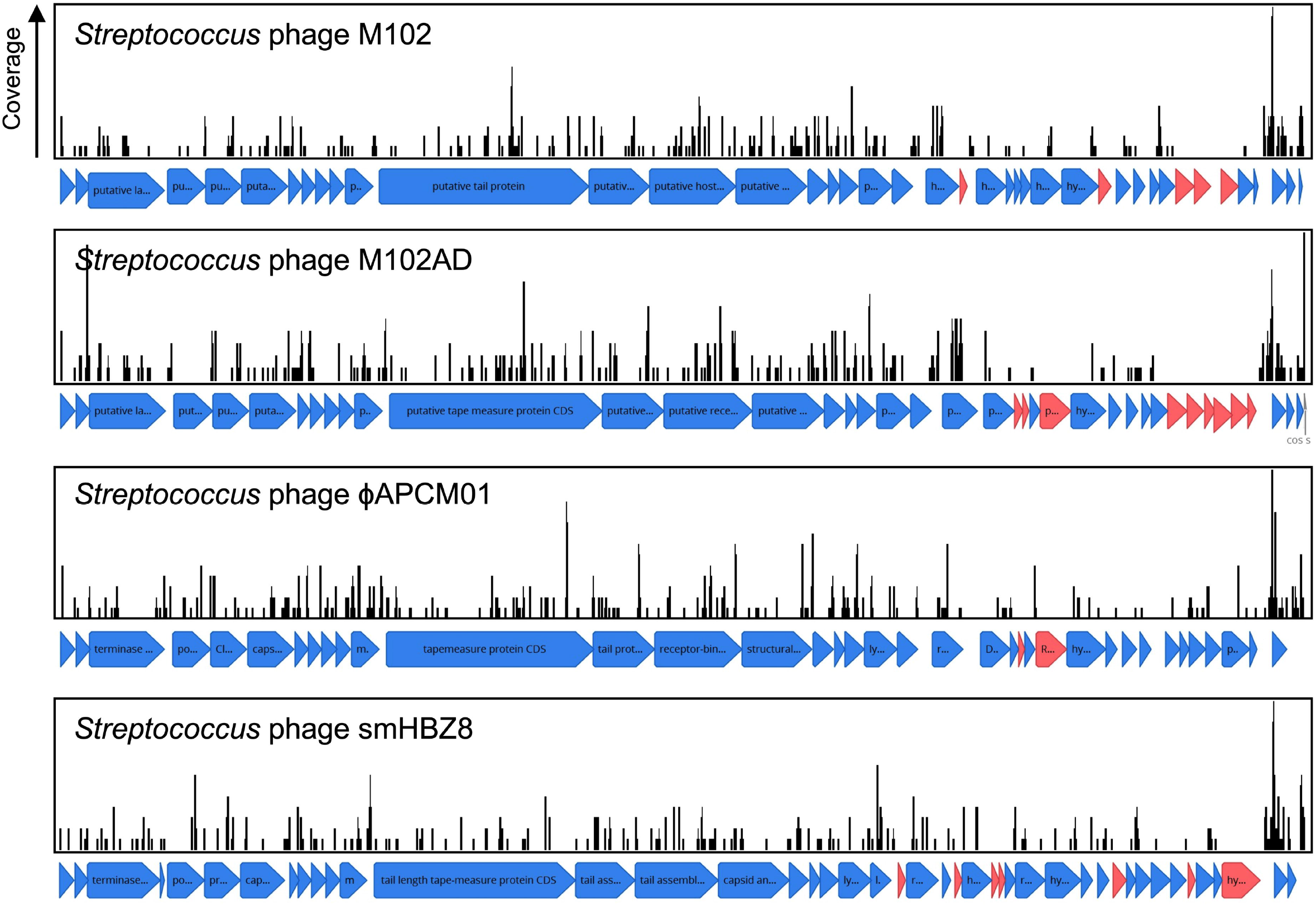
CRISPR spacer acquisition against *S. mutans* bacteriophage. CRISPR spacers were blasted against viral sequences downloaded from NCBI. CRISPR spacers mapped to four *S. mutans* infecting phage, M102, M102AD, □APCM01 and smHBZ8. These are the only *S. mutans* phage that have been genome sequenced and all belong to the *Siphoviridae* family of double-stranded DNA viruses. CRISPR spacers that mapped to phage sequences are shown as black lines, with the height of these lines indicating the total number of spacers that mapped to that specific region. Open reading frames for which no CRISPR spacers have been acquired against are colored red, and are more frequently located in a variable region of the phage genomes.

Isolation of anti-*S. mutans* phage is rare. For example, □APCM01 was the only *S. mutans* phage isolated from 85 saliva samples [33] and smHBZ8 was the single *S. mutans* phage isolated from 254 samples (diverse sources; saliva, dental sewage, cariogenic dentin, extracted teeth, and dental plaque) [31]. To understand how important CRISPR-Cas is in explaining the low frequency of infective phage isolation we calculated the frequency of phage immunity spacers across our genome database. In total, 64% of *S. mutans* strains that carry CRISPR-Cas systems have phage targeting spacers (against M102AD or M102 or □APCM01). The high frequency of CRISPR-Cas systems and anti-phage spacers would likely limit the efficacy of phage therapy in *S. mutans*.

#### Plasmids

CRISPR spacers are also naturally acquired against plasmid DNA [14, 35]. Acquisition of spacers against plasmids was very infrequently identified for *S. mutans*. In a database containing ∼12,000 sequenced plasmids [19] only two spacers mapped. A spacer from smu333 mapped to pLF25067 carried by *Lactobacillus fermentum* [36] and a spacer from smu408 mapped to pSZ4 from *Staphylococcus warneri* [37]. One strain, smu441, carries a spacer that maps (88% coverage) against a small 5.6 kbp ‘cryptic’ plasmid that is carried by ∼5% of *S. mutans* strains [20, 21].

#### Integrative conjugative plasmids

Integrative and conjugative elements (ICEs; also known as conjugative transposons) are another type of mobile genetic element that can transfer between bacterial species and strains [38]. Sequence data for known ICEs was downloaded from ICEberg and used to check for CRISPR spacer acquisition against ICEs. In total, *S. mutans* has acquired fifty-eight spacers against ICEs. All but one of these spacers matches against TnSmu1, an uncharacterized putative ICE which is related to Tn916 but with significant modifications [39]. In the model *S. mutans* strain UA159 TnSmu1 is a large 20 kbp region comprising of over twenty predicted genes [39]. Most of the genes in this region are hypothetical but some are predicted to have integrative and conjugative functions (*e*.*g*. type IV secretion system components). When aligned to this region the CRISPR spacers qualitatively appear to have been generated to target the integrase and secretion system components instead of hypothetical regions (Figure 3). The functionality of TnSmu1 and whether it behaves like an ICE is unknown. However, acquisition of CRISPR spacers by strains of *S. mutans* could indicate that it may be horizontally acquired and strains are actively trying to avoid acquisition of the element. Certain strains of *S. mutans* may also have been repeatedly attacked by TnSmu1 as the isolate Smu179 has acquired five spacers that align to TnSmu1. In *Pseudomonas aeruginosa* CRISPR spacers often target ICE or conjugative transfer genes and this is associated with a lower prevalence of these systems in strains harboring CRISPR-Cas systems [13]. Prevention of the acquisition of these elements likely arises because of fitness costs associated with them. Our analysis of CRISPR-mediated immunity of ICEs is likely conservative as description and characterization of these elements is limited in *S. mutans*.

**Figure 3.**
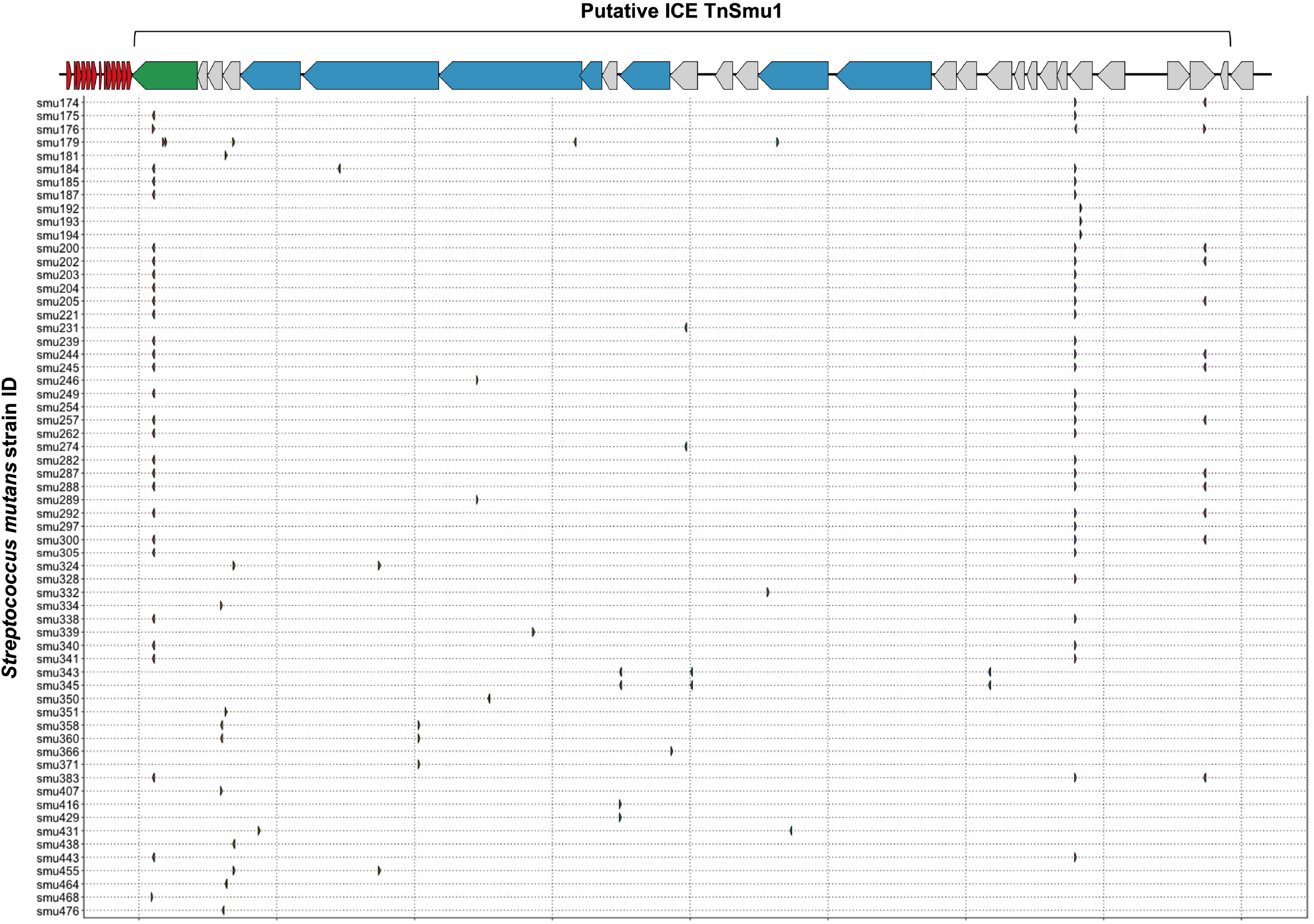
CRISPR spacer acquisition against the putative integrative and conjugative element TnSmu1. CRISPR spacers were blasted against ICE sequences downloaded from ICEberg. Multiple CRISPR spacers mapped to a putative *S. mutans* ICE known as TnSmu1. This element is approximately 20 Kbp and consists of twenty-eight genes. Most of the genes are hypothetical (grey) but some are annotated as putative T4SS components (blue) and one gene is annotated as an integrase (green). CRISPR spacers that mapped to this region are shown as small arrows, with a row for each *S. mutans* strain that has acquired spacers against the putative ICE.

### *S. mutans* self- and strain-targeting CRISPR spacers

With the observation that *S. mutans* has gained immunity against inter-species transfer of mobile genetic elements we next wanted to explore if CRISPR spacers have been acquired more generally against *S. mutans* strain DNA. To do this, we searched for CRISPR spacer hits within genes encoded by *S. mutans* across all the genomes included in this study. A total of sixty-three annotated genes are the target of CRISPR spacers (Table S2). In addition, six other genes with functional predictions (Table S2) are also the target of CRISPR spacers. Lastly, *S. mutans* hypothetical proteins were also matched to *S. mutans* CRISPR spacer sequences. From this set of genes (excluding hypotheticals), 22 reside in the core genome (>99% of genomes), 4 are soft core genes (95-99% of genomes), 17 are shell genes (15-95% of genomes), and 20 are cloud genes (<15% of genomes) (Table S2). CRISPR targets against *S. mutans* core genes are over-represented given that core genes only account for 5% of the *S. mutans* pangenome. This is compared with cloud genes which make up 85% of the pangenome. Notable *S. mutans* gene CRISPR targets include the *clp* system [40], the *gtfC* glucosyltransferase [41], the cell surface antigen *spaP* [42], carbohydrate utilization systems (i.e. *fruA* and *lacF*) [43, 44] and several genes involved in DNA replication, repair or modification (i.e. *dpnM, parE, radD, recF, repA, smc, ssb* and *topB*). Several of the CRISPR spacers that target the *S. mutans* pangenome are likely targeting mobile DNA elements. For example, spacers against tyrosine recombinases (i.e. *xerC* and *xerD*), conjugal transfer protein *traG*, and the Int-Tn transposase from the ICE Tn916. To visualize the extent of CRISPR spacer acquisition against *S. mutans* genomes, spacers were mapped to two strains, UA159 and NN2025 (Figure 4A and B). The number of spacers that aligned within 10 kb regions varied, with notable hotspots against the genomes of both UA159 and NG8. These hotspot regions were TnSmu1 and repeat regions within the promoters of *clpP* (protein homeostasis), *rexB* (putative exonuclease) and a hypothetical protein.

**Figure 4.**
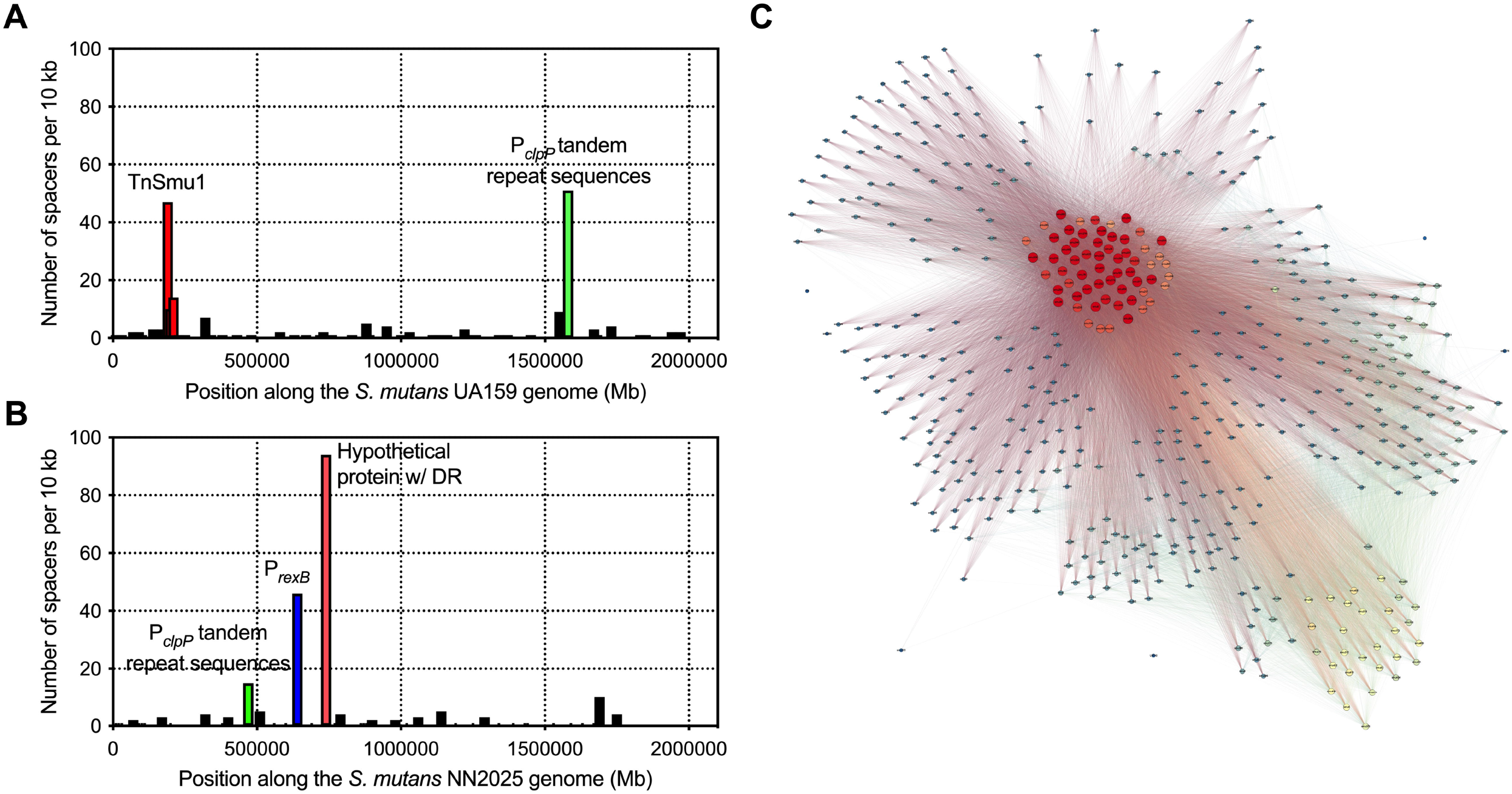
Acquisition of CRISPR spacers against *S. mutans* genes. To visualize CRISPR spacer acquisition against *S. mutans* genomes, spacers were mapped to two strains, UA159 (A) and NN2025 (B). Each graph shows the number of spacers that aligned within each 10 Kbp region of a *S. mutans* genome. There are notable hotspots that are highlighted with red, green and blue bars. A network diagram was constructed using Gephi (http://www.gephi.org) to visualize *S. mutans* strains that interact with other *S. mutans* strains on the basis of CRISPR spacer acquisition (C). Each node (circle) is a single *S. mutans* strain and a line between strains indicates a connection based on a CRISPR target being present on one of those strains. Strains with increased connections (targeting more strains with CRISPR spacers) are colored on a scale, with red nodes depicting strains with the most connections (CRISPR spacers targeting other *S. mutans* strains).

Inter-species CRISPR targeting was also visualized using network analysis (Figure 4C). This analysis showed that a minority of *S. mutans* strains (ca. 60 strains; red nodes in Figure 4C) were accounting for the majority of CRISPR spacer connections between *S. mutans* strains. These strains carry CRISPR spacers against highly conserved (core) *S. mutans* genes (Table S2). We reasoned that this group of highly connected strains could actually be carrying self-targeting CRISPR spacers against highly conserved genes. Analysis of self-targeting revealed that ninety-four *S. mutans* strains carry self-targeting CRISPR spacers (Table S3) and most of the highly connected nodes in the network analysis are strains that self-target. All of these strains contain the Cas genes required for functional CRISPR-Cas systems but it is unknown if these systems are inactivated through point mutations. Acquisition of self-targeting spacers would likely be detrimental and create pressure for inactivation of the CRISPR-Cas systems. Six out of the ninety-four self-targeting strains contain anti-CRISPR (Acr) proteins that could also account for a loss of functionality (Table S3). In summary, *S. mutans* strains carry spacers that target *S. mutans* genes although the relative amount of intra-species-targeting versus only self-targeting is currently unknown.

### Limited impact of CRISPR-Cas systems on *S. mutans* genome size and GC content

Analysis of *S. mutans* CRISPR spacers has shown potential immunity acquisition against phage, plasmid, ICE, and *S. mutans* DNA. Acquisition of spacers against these types of DNA should act to limit the rate of HGT in *S. mutans*. The association between CRISPR-Cas and its impact on HGT in bacteria is not clear. For example, *P. aeruginosa* genomes with CRISPR-Cas are significantly smaller than those lacking CRISPR-Cas which suggests that CRISPR-Cas impedes HGT [13]. However, a similar trend was not found for *Staphylococcus aureus* or *Acinetobacter baumannii* [45]. Here, we are using CRISPR spacer load as a measure of CRISPR-Cas activity. It is unfeasible to test for genomic differences between strains with and without CRISPR-Cas systems because the majority of *S. mutans* strains have at least one CRISPR-Cas system. CRISPR spacer load, the number of CRISPR spacers per genome, has been used as a measure of CRISPR activity by others [15]. For our analysis, *S. mutans* strains were grouped by the total number of spacers they have acquired and for each group the median size of the genome was computed (Figure 5). We found that increased CRISPR spacers loads were associated with marginally increased genome sizes. Strains with 1-20 CRISPR spacers had a median genome size of 2.01 Mbp and strains with 40-60 CRISPR spacers had a median genome size of 2.05 Mbp (two-tailed *t*-test *p* < 0.01). Our analysis shows that increased CRISPR spacer acquisition does not lead to reduced genome size in *S. mutans*.

**Figure 5.**
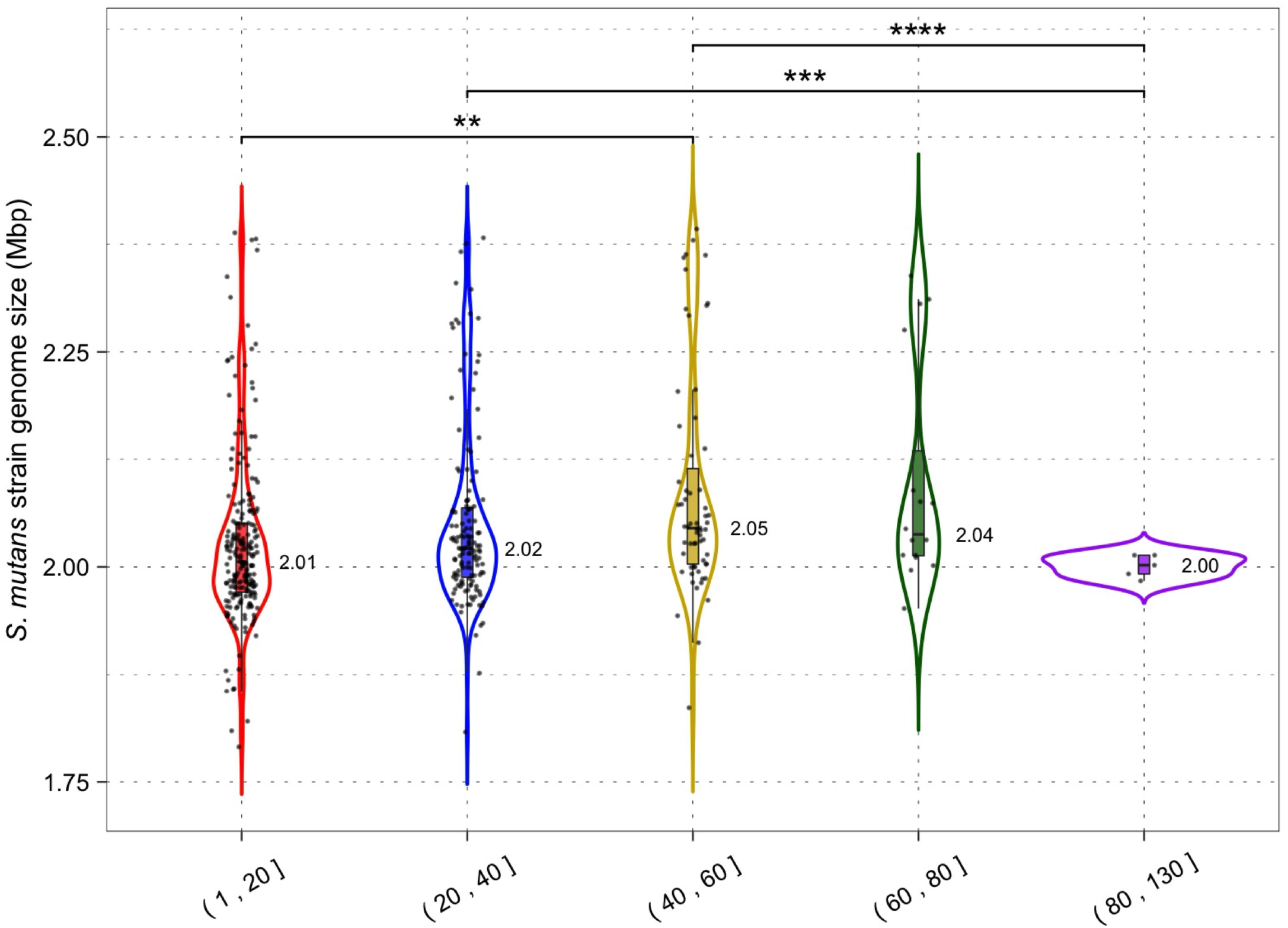
CRISPR activity and genome size. Genomes were grouped according to the number of CRISPR spacers present (1-20, 21-40, 41-60, 61-80 and 81+) and a violin plot was generated for each group. Median genome size (Mbp) is shown next to each violin plot; each dot is a single *S. mutans* strain/genome. Statistical significance between groups was calculated using a two-tailed *t*-test (** = *p* < 0.01; *** = *p* < 0.001).

By limiting HGT CRISPR-Cas could also affect the GC content of host genomes by reducing the acquisition of higher or lower GC foreign DNA. The average GC content across all *S. mutans* CRISPR spacers was 38.2% (Fig. 1D) which may indicate that on average foreign DNA elements have higher GC content than the average *S. mutans* genome (36.8%). Phages that infect *S. mutans*, M102AD, M102, and SMHBZ8 have GC content of 39.6%, 39.21% and 38.8% respectively [31, 34, 46]. The GC content of *S. mutans* genomes was consistent for strains with 1-20 (36.80%), 20-40 (36.82%) and 40-60 (36.79%) CRISPR spacers (Figure 6). A significant difference in GC content was observed for strains with 60-80 (36.68%) CRISPR spacers compared with the 1-20 and 20-40 groupings (two-tailed *t*-test *p* < 0.05). Although only nominally lower than the other groups, this might indicate a reduction in the acquisition of higher GC foreign DNA among strains with 60-80 CRISPR spacers.

**Figure 6.**
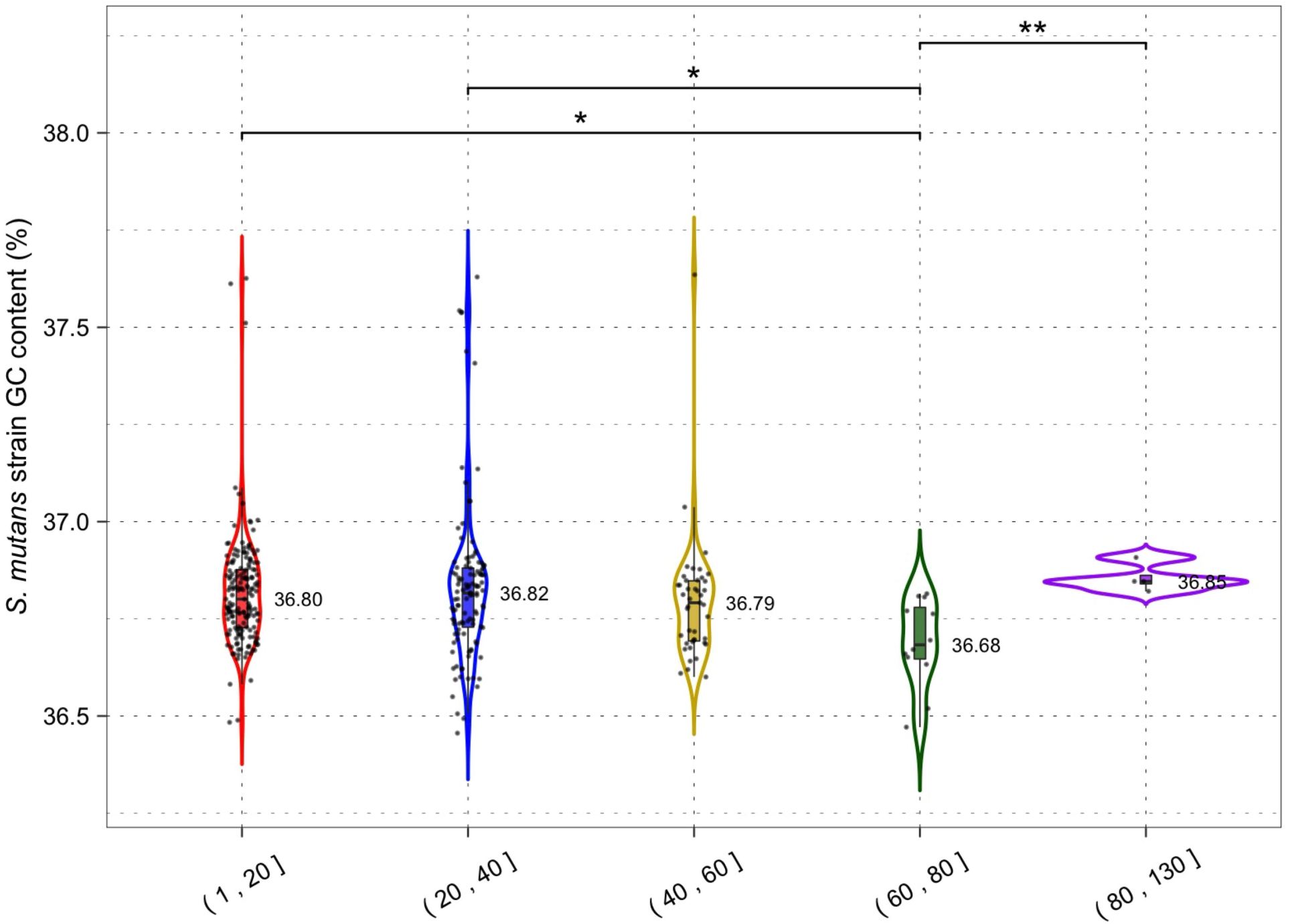
CRISPR activity and GC content. Genomes were grouped according to the number of CRISPR spacers present (1-20, 21-40, 41-60, 61-80 and 81+) and a violin plot was generated for each group. Median GC content (percentage GC) is shown next to each violin plot; each dot is a single *S. mutans* strain/genome. Statistical significance between groups was calculated using a two-tailed *t*-test (* = *p* < 0.05; ** = *p* < 0.01).

### CRISPR-Cas systems do not restrict the size of the *S. mutans* pangenome

To gain a greater insight into the impact of CRISPR-Cas systems on HGT we wanted to examine how CRISPR spacer load alters the size and distribution of the *S. mutans* pangenome. In our analysis including 335 *S. mutans* genomes, the pangenome size was calculated to include 15,062 genes. There were 822 genes in the core genome (>99% of genomes), 496 soft core genes (95-99% of genomes), 982 shell genes (15-95% of genomes), and 12,762 cloud genes (<15% of genomes). Next, pangenome analysis was conducted on two groups, one with a low CRISPR spacer load (1-20 spacers, average 9.36; 163 genomes) and another with a high CRISPR spacer load (21-129 spacers, average 37.93; 169 genomes). For both groups we generated pie charts which show the percentage of genes located in the different gene clusters (core; soft core; shell; cloud) (Figure 7). The low and high CRISPR spacer load groups have a similar percentage of genes located within the cloud genes gene cluster. Cloud genes are those that are most likely acquired via HGT. By this measure there is no limiting of HGT by CRISPR-Cas systems for *S. mutans* strains with either low or high CRISPR spacer loads.

**Figure 7.**
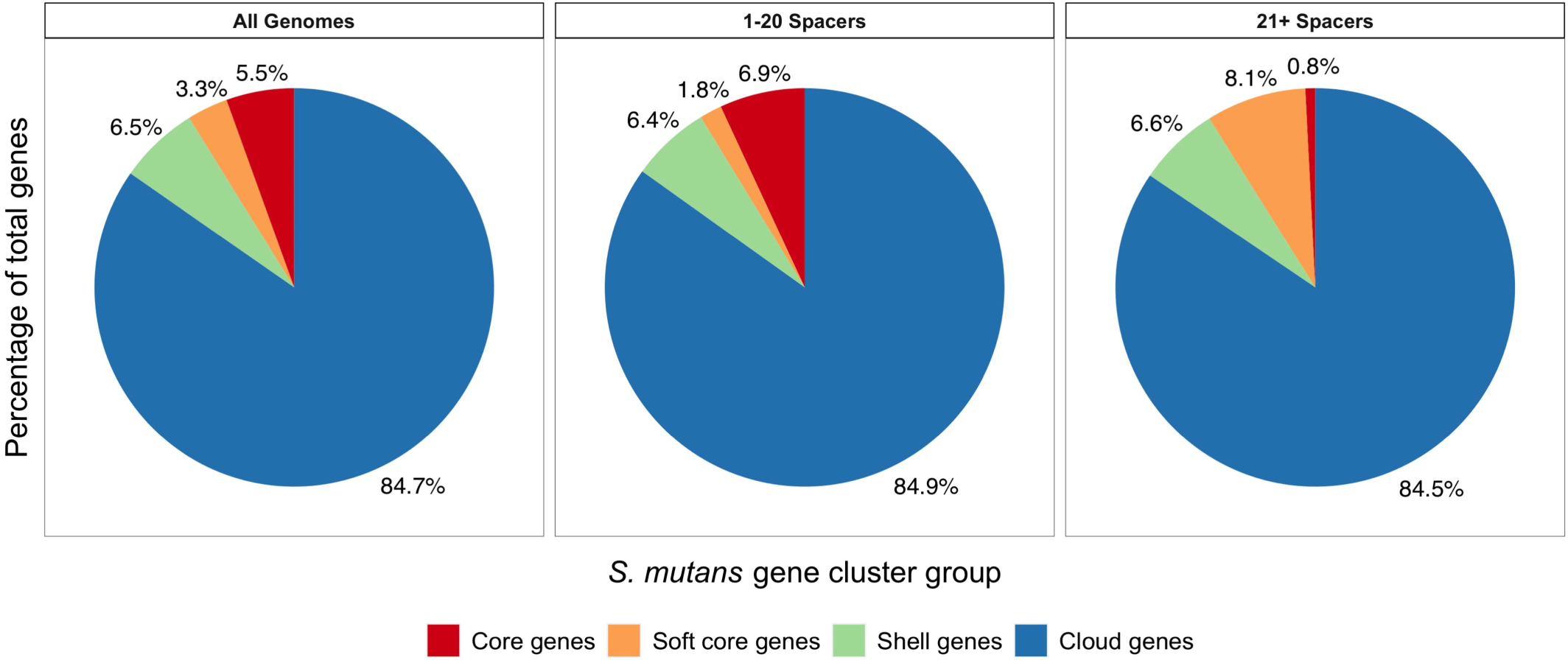
Pangenome analysis comparing low and high CRISPR spacer load *S. mutans* strains. Pangenome analysis was conducted on three groups (1) all *S. mutans* genomes, (2) a group with low CRISPR spacer load (1-20 spacers, average 9.36; 163 genomes) and (3) a group with a high CRISPR spacer load (21-129 spacers, average 37.93; 169 genomes). For the three groups pie charts were generated which show the percentage of genes located in the different gene clusters (core, red; soft core, orange; shell, green; cloud, blue). Cloud genes (blue) are most likely acquired via HGT.

## Conclusion

Here, we have generated a comprehensive ‘memory’ of DNA that has attacked a diverse panel of *S. mutans* isolates. Although there is considerable ‘dark matter’ (undetected CRISPR spacer sequences), phage are the main target of *S. mutans* spacers. Isolates have also generated immunity against mobile DNA elements such as plasmids and ICEs. There may also be considerable immunity generated against bacterial DNA, although the relative contribution of self-targeting versus bona fide intra- or inter-species targeting needs to be investigated further. Collectively, CRISPR-Cas systems and acquired spacers are abundant across all *S. mutans* strains. While there is clear evidence that these systems have acquired immunity against foreign DNA, there appears to be minimal impact on HGT constraints on a species-level. There was little or minimal impact on genome size, GC content and ‘openness’ of the pangenome. There are several possible explanations for why there is a minimal impact on HGT and these include (1) the systems have specificity for phage infection, (2) self-targeting has led to a loss of activity, (3) the systems are highly regulated to only acquire immunity under certain conditions, (4) competence acquired DNA is single stranded and the CRISPR-Cas systems are less active against ssDNA, and (5) spacer load may be an inaccurate measure of CRISPR-Cas activity. A deeper understanding of the impact of these systems on the biology and evolution of *S. mutans* will likely require functional studies and/or model systems for interrogating spacer acquisition.

## Acknowledgements

We would like to thank Vince Richards (Clemson University) for providing us with the *S. mutans* genome files associated with this study.

## Author contributions

A.R.W. and R.C.S. were involved in the investigation, data curation and revised the manuscript; A.R.W. performed the formal analysis and software development (implementation of code and supporting algorithms); R.C.S. was involved with funding acquisition, conceptualization of the study and drafted the manuscript.

