## Supplemental Material for "Investigating CRISPR spacer targets and their impact on genomic diversification of *Streptococcus mutans*"

Alejandro R. Walker[1] and Robert C. Shields[2]*
[1] Department of Oral Biology, University of Florida, Gainesville, Florida, USA
[2] Department of Biological Sciences, Arkansas State University, Jonesboro, Arkansas, USA

### Supplementary Table S1. Bacteriophage targeted by *S. mutans* CRISPR spacers.

| Phage strain | CRISPR spacer sequence | Target | Host |
|---|---|---|---|
| **Enterococcus phage vB_EfaS_IME197** | TATCTAACCTATCCTTACCTAACCTTACCT | replication initiation protein | *E. faecalis* |
| **Enterococcus phage vB_EfaS_IME197** | ATCTAACCTATCCTTACCTAACCTTACCTG | replication initiation protein | *E. faecalis* |
| **Lactobacillus phage LF1** | TCTAACCTATCCTTACCTAACCTTACCTG | replication initiation protein | *L. fermentum* |
| **Staphylococcus phage CNPH82** | TGTTTACCGTCATAATAGCAAGGTAAGAT | conserved phage protein | *S. epidermidis* |
| **Staphylococcus phage CNPx** | TGTTTACCGTCATAATAGCAAGGTAAGAT | conserved phage protein | *S. epidermidis* |
| **Staphylococcus phage PH15** | TGTTTACCGTCATAATAGCAAGGTAAGAT | conserved phage protein | *S. epidermidis* |
| **Staphylococcus phage vB_SepiS-philPLA5** | TGTTTACCGTCATAATAGCAAGGTAAGAT | conserved phage protein | *S. epidermidis* |
| **Staphylococcus phage vB_SepiS-philPLA7** | TGTTTACCGTCATAATAGCAAGGTAAGAT | conserved phage protein | *S. epidermidis* |
| **Streptococcus virus Sfi21** | AGAGAACTCGAAGAAAATACCGATAAGACA | orf123 | *S. thermophilus* |
| **Streptococcus phage 7201** | TGGGTGCTAAAGGTGATGACTATCGTTTCA | orf33 | *S. thermophilus* |
| **Streptococcus phage A25** | ACTTGCATATACACATTTTGTTCAACATCA | portal protein | *S. pyogenes* |
| **Streptococcus phage APCM01** | multiple spacers | | *S. mutans* |
| **Streptococcus phage M102** | multiple spacers | | *S. mutans* |
| **Streptococcus phage M102AD** | multiple spacers | | *S. mutans* |
| **Streptococcus phage smHBZ8** | multiple spacers | | *S. mutans* |
| **Streptococcus phage phiNJ2** | ACTTGCATATACACATTTTGTTCAACATCA | portal protein | *S. suis* |
| **Streptococcus phage T12** | TTATCAAGTTGTCCATACTGAGCTTTCATAATT | putative methyltransferase | *S. pyogenes* |
| **Streptococcus phage T12** | AATTATGAAAGCTCAGTATGGACAACTTGATAA | putative methyltransferase | *S. pyogenes* |
| **Streptococcus phage T12** | TATCAAGTTGTCCATACTGAGCTTTCATAATT | putative methyltransferase | *S. pyogenes* |

### Supplementary Table S2. *S. mutans* genes targeted by *S. mutans* CRISPR spacers.

| Gene | Product | Core Genome | Evidence Self Target-targeting |
|---|---|---|---|
| ***ackA*** | Acetate kinase | Core | Yes |
| ***adeP*** | Adenine permease AdeP | Cloud | |
| ***aroA*** | 3-phosphoshikimate 1-carboxyvinyltransferase | Core | Yes |
| ***atpD*** | ATP synthase subunit beta | Core | |
| ***cbl*** | HTH-type transcriptional regulator cbl | Core | Yes |
| ***clpC*** | ATP-dependent Clp protease ATP-binding subunit ClpC | Core | |
| ***clpL*** | ATP-dependent Clp protease ATP-binding subunit ClpL | Cloud | |

| | | | |
|---|---|---|---|
| ***clpP1*** | ATP-dependent Clp protease proteolytic subunit 1 | Cloud | |
| ***clpX*** | ATP-dependent Clp protease ATP-binding subunit ClpX | Core | Yes |
| ***coaD*** | Phosphopantetheine adenylyltransferase | Core | Yes |
| ***comEC*** | ComE operon protein 3 | Shell | Yes |
| ***dpnM*** | Modification methylase DpnIIA | Cloud | |
| ***ebh*** | Extracellular matrix-binding protein ebh | Cloud | Yes |
| ***esaA*** | ESAT-6 secretion accessory factor EsaA | Cloud | Yes |
| ***fccA*** | Fumarate reductase flavoprotein subunit | Shell | |
| ***fhaB*** | Filamentous hemagglutinin | Cloud | Yes |
| ***fieF*** | Ferrous-iron efflux pump FieF | Core | |
| ***fruA*** | Fructan beta-fructosidase | Core | |
| ***glnR*** | HTH-type transcriptional regulator GlnR | Core | Yes |
| ***grsB*** | Gramicidin S synthase 2 | Shell | Yes |
| ***gtfC*** | Glucosyltransferase-SI | Shell | Yes |
| ***hcaR*** | Hca operon transcriptional activator HcaR | Cloud | |
| ***hin*** | DNA-invertase hin | Cloud | |
| ***hisS*** | Histidine--tRNA ligase | Core | Yes |
| ***Int-Tn*** | Transposase from transposon Tn916 | Shell | |
| ***lacF*** | Lactose transport system permease protein LacF | Core | |
| ***lgrD*** | Linear gramicidin synthase subunit D | Shell | |
| ***lpxD*** | UDP-3-O-(3-hydroxymyristoyl)glucosamine N-acyltransferase | Cloud | |
| ***ltaS2*** | Lipoteichoic acid synthase 2 | Core | Yes |
| ***ltrA*** | Group II intron-encoded protein LtrA | Cloud | |
| ***mhbT*** | 3-hydroxybenzoate transporter MhbT | Shell | Yes |
| ***mleA*** | Malolactic enzyme | Shell | Yes |
| ***mrp*** | Iron-sulfur cluster carrier protein | Core | Yes |
| ***nfr1*** | NADH-dependent flavin reductase subunit 1 | Soft Core | Yes |
| ***nrdG*** | Anaerobic ribonucleoside-triphosphate reductase-activating protein | Cloud | |
| ***parE*** | DNA topoisomerase 4 subunit B | Core | Yes |
| ***pbuG*** | Guanine/hypoxanthine permease PbuG | Core | Yes |
| ***plsX*** | Phosphate acyltransferase | Core | Yes |
| ***ppsB*** | Plipastatin synthase subunit B | Shell | Yes |
| ***ppsC*** | Plipastatin synthase subunit C | Shell | Yes |
| ***pyrG*** | CTP synthase | Core | Yes |
| ***radD*** | Putative DNA repair helicase RadD | Cloud | Yes |
| ***recF*** | DNA replication and repair protein RecF | Core | |
| ***repA*** | Regulatory protein RepA | Cloud | |
| ***rlmCD*** | 23S rRNA (uracil-C(5))-methyltransferase RlmCD | Soft Core | |
| ***rnmV*** | Ribonuclease M5 | Soft Core | Yes |

| | | | |
|---|---|---|---|
| ***secA*** | Protein translocase subunit SecA | Core | Yes |
| ***smc*** | Chromosome partition protein Smc | Soft Core | |
| ***soj*** | Sporulation initiation inhibitor protein Soj | Cloud | |
| ***spaP*** | Cell surface antigen I/II | Shell | Yes |
| ***ssaA*** | Staphylococcal secretory antigen SsaA | Cloud | |
| ***ssb*** | Single-stranded DNA-binding protein | Core | |
| ***ssrA*** | transfer-messenger RNA%2C SsrA | | Yes |
| ***tilS*** | tRNA(Ile)-lysidine synthase | Core | |
| ***topB*** | DNA topoisomerase 3 | Cloud | |
| ***traG*** | Conjugal transfer protein TraG | Shell | |
| ***ubiE*** | Ubiquinone/menaquinone biosynthesis C-methyltransferase UbiE | Shell | |
| ***urdA*** | Urocanate reductase | Cloud | |
| ***valS*** | Valine--tRNA ligase | Shell | Yes |
| ***xerC*** | Tyrosine recombinase XerC | Shell | |
| ***xerD*** | Tyrosine recombinase XerD | Shell | |
| ***xre*** | HTH-type transcriptional regulator Xre | Cloud | |
| ***yhdJ*** | DNA adenine methyltransferase YhdJ | Cloud | |
| **(blank)** | hypothetical protein | n/a | Variable |
| **(blank)** | N-acetylmuramoyl-L-alanine amidase domain-containing protein | Shell | Yes |
| **(blank)** | putative ABC transporter ATP-binding protein | | |
| **(blank)** | putative cation efflux system protein | | Yes |
| **(blank)** | Putative multidrug export ATP-binding/permease protein | | Yes |
| **(blank)** | putative multidrug-efflux transporter | | |
| **(blank)** | Ribonuclease | | |

### Supplementary Table S3. *S. mutans* strains that carry self-targeting CRISPR spacers.

| Strain | Self-target gene | Self-target product | Contains Acr Protein |
|---|---|---|---|
| **smu125** | nfr1_2 | NADH-dependent flavin reductase subunit 1 | |
| **smu173** | | hypothetical protein | Yes |
| **smu174** | ssrA | transfer-messenger RNA%2C SsrA | |
| **smu179** | gtfC_6 | Glucosyltransferase-SI | |
| **smu179** | gtfC_6 | Glucosyltransferase-SI | |
| **smu179** | gtfC_6 | Glucosyltransferase-SI | |
| **smu179** | lacF_2 | Lactose transport system permease protein LacF | |
| **smu179** | | hypothetical protein | |
| **smu179** | | hypothetical protein | |
| **smu181** | ackA | Acetate kinase | |
| **smu184** | valS | Valine--tRNA ligase | |
| **smu185** | valS_1 | Valine--tRNA ligase | |
| **smu185** | valS_2 | Valine--tRNA ligase | |

| | | |
|---|---|---|
| **smu187** | valS | Valine--tRNA ligase |
| **smu192** | lgrD_2 | Linear gramicidin synthase subunit D |
| **smu193** | lgrD_2 | Linear gramicidin synthase subunit D |
| **smu194** | lgrD_2 | Linear gramicidin synthase subunit D |
| **smu201** | | hypothetical protein |
| **smu209** | nfr1_1 | NADH-dependent flavin reductase subunit 1 |
| **smu216** | secA | Protein translocase subunit SecA |
| **smu217** | gtfC_1 | Glucosyltransferase-SI |
| **smu224** | rnmV | Ribonuclease M5 |
| **smu225** | rnmV_1 | Ribonuclease M5 |
| **smu238** | mleA | Malolactic enzyme |
| **smu240** | secA | Protein translocase subunit SecA |
| **smu251** | aroA | 3-phosphoshikimate 1-carboxyvinyltransferase |
| **smu254** | | hypothetical protein |
| **smu262** | cbl | HTH-type transcriptional regulator cbl |
| **smu263** | rnmV | Ribonuclease M5 |
| **smu267** | rnmV | Ribonuclease M5 |
| **smu268** | rnmV_1 | Ribonuclease M5 |
| **smu281** | mleA | Malolactic enzyme |
| **smu283** | secA | Protein translocase subunit SecA |
| **smu294** | aroA | 3-phosphoshikimate 1-carboxyvinyltransferase |
| **smu297** | | hypothetical protein |
| **smu305** | cbl | HTH-type transcriptional regulator cbl |
| **smu306** | rnmV | Ribonuclease M5 |
| **smu315** | mleA | Malolactic enzyme |
| **smu318** | | hypothetical protein |
| **smu320** | | hypothetical protein |
| **smu324** | grsB_1 | Gramicidin S synthase 2 |
| **smu324** | rlmCD_2 | 23S rRNA (uracil-C(5))-methyltransferase RlmCD |
| **smu324** | rlmCD_2 | 23S rRNA (uracil-C(5))-methyltransferase RlmCD |
| **smu327** | hisS | Histidine--tRNA ligase |
| **smu327** | | hypothetical protein |
| **smu327** | | hypothetical protein |
| **smu327** | | hypothetical protein |
| **smu327** | | hypothetical protein |
| **smu329** | hisS | Histidine--tRNA ligase |
| **smu332** | | hypothetical protein |
| **smu333** | | hypothetical protein |
| **smu335** | | putative ABC transporter ATP-binding protein |
| **smu336** | radD | Putative DNA repair helicase RadD |
| **smu337** | spaP_2 | Cell surface antigen I/II |
| **smu337** | | hypothetical protein |
| **smu338** | cbl | HTH-type transcriptional regulator cbl |
| **smu34** | parE | DNA topoisomerase 4 subunit B |

| | | | |
|---|---|---|---|
| **smu340** | cbl | HTH-type transcriptional regulator cbl | |
| **smu341** | cbl | HTH-type transcriptional regulator cbl | |
| **smu343** | fccA_2 | Fumarate reductase flavoprotein subunit | |
| **smu344** | | hypothetical protein | |
| **smu345** | fccA_2 | Fumarate reductase flavoprotein subunit | |
| **smu349** | | hypothetical protein | |
| **smu362** | fruA_2 | Fructan beta-fructosidase | |
| **smu371** | pbuG | Guanine/hypoxanthine permease PbuG | |
| **smu372** | fhaB | Filamentous hemagglutinin | |
| **smu372** | fhaB | Filamentous hemagglutinin | |
| **smu375** | esaA_2 | ESAT-6 secretion accessory factor EsaA | Yes |
| **smu375** | smc_6 | Chromosome partition protein Smc | Yes |
| **smu376** | esaA_2 | ESAT-6 secretion accessory factor EsaA | Yes |
| **smu376** | smc_6 | Chromosome partition protein Smc | Yes |
| **smu378** | coaD | Phosphopantetheine adenylyltransferase | |
| **smu378** | ppsB | Plipastatin synthase subunit B | |
| **smu382** | atpD | ATP synthase subunit beta | |
| **smu383** | | hypothetical protein | |
| **smu39** | pyrG | CTP synthase | |
| **smu390** | ppsC | Plipastatin synthase subunit C | |
| **smu391** | ppsC | Plipastatin synthase subunit C | |
| **smu395** | | hypothetical protein | |
| **smu401** | ebh | Extracellular matrix-binding protein ebh | |
| **smu401** | | hypothetical protein | |
| **smu409** | smc_2 | Chromosome partition protein Smc | |
| **smu410** | | hypothetical protein | |
| **smu411** | | hypothetical protein | |
| **smu411** | | hypothetical protein | |
| **smu412** | nfr1_2 | NADH-dependent flavin reductase subunit 1 | |
| **smu414** | | hypothetical protein | Yes |
| **smu416** | comEC | ComE operon protein 3 | |
| **smu416** | | hypothetical protein | |
| **smu419** | | hypothetical protein | Yes |
| **smu422** | mhbT | 3-hydroxybenzoate transporter MhbT | |
| **smu422** | mhbT | 3-hydroxybenzoate transporter MhbT | |
| **smu423** | clpX | ATP-dependent Clp protease ATP-binding subunit ClpX | |
| **smu423** | ppsC | Plipastatin synthase subunit C | |
| **smu426** | | hypothetical protein | Yes |
| **smu431** | ltaS2 | Lipoteichoic acid synthase 2 | |
| **smu445** | glnR | HTH-type transcriptional regulator GlnR | |
| **smu445** | lgrD_2 | Linear gramicidin synthase subunit D | |
| **smu445** | | hypothetical protein | |
| **smu454** | | hypothetical protein | |
| **smu472** | mrp | Iron-sulfur cluster carrier protein | |

| | | |
|---|---|---|
| **smu6** | | hypothetical protein |
| **smu6** | | putative cation efflux system protein |
| **smu76** | plsX | Phosphate acyltransferase |
